# Scalable DNA Feature Generation and Transcription Factor Binding Prediction via Deep Surrogate Models

**DOI:** 10.1101/2024.12.06.626709

**Authors:** Anowarul Kabir, Toki Tahmid Inan, Kim Rasmussen, Amarda Shehu, Anny Usheva, Alan Bishop, Boian Alexandrov, Manish Bhattarai

## Abstract

Simulating DNA breathing dynamics, for instance Extended Peyrard-Bishop-Dauxois (EPBD) model, across the entire human genome using traditional biophysical methods like pyDNA-EPBD is computationally prohibitive due to intensive techniques such as Markov Chain Monte Carlo (MCMC) and Langevin dynamics. To overcome this limitation, we propose a deep surrogate generative model utilizing a conditional Denoising Diffusion Probabilistic Model (DDPM) trained on DNA sequence-EPBD feature pairs. This surrogate model efficiently generates high-fidelity DNA breathing features conditioned on DNA sequences, reducing computational time from months to hours–a speedup of over 1000 times. By integrating these features into the EPBDxDNABERT-2 model, we enhance the accuracy of transcription factor (TF) binding site predictions. Experiments demonstrate that the surrogate-generated features perform comparably to those obtained from the original EPBD framework, validating the model’s efficacy and fidelity. This advancement enables real-time, genome-wide analyses, significantly accelerating genomic research and offering powerful tools for disease understanding and therapeutic development.

## 1 Introduction

Accurate modeling of DNA dynamics is essential for understanding genetic regulation and disease mechanisms. DNA breathing—the spontaneous local opening and closing of the double-helix structure—plays a significant role in gene expression regulation by influencing transcription factor (TF) binding and gene expression (Nowak-Lovato et al. 2013a; Alexandrov et al. 2012, 2010). Traditional biophysical simulation methods, such as Markov Chain Monte Carlo (MCMC) and Langevin dynamics employed in frameworks like pyDNA-EPBD (Kabir et al. 2023a,b; Choi et al. 2008), offer high-fidelity modeling of DNA breathing dynamics. However, simulating the entire human genome using these methods is computationally intensive and time-consuming, often requiring months (Alexandrov et al. 2010), which limits large-scale genomic studies.

To address these challenges, we propose a deep surrogate generative modeling approach that leverages a conditional Denoising Diffusion Probabilistic Model (DDPM) (Ho, Jain, and Abbeel 2020) to efficiently generate DNA breathing features conditioned on DNA sequences. By training the surrogate model on a fraction of sequence-feature pairs generated using existing frameworks like pyDNA-EPBD, we enable the model to learn complex relationships between DNA sequences and their corresponding biophysical properties. Once trained, this model can generate DNA breathing features for the entire human genome within few days, a task previously deemed computationally infeasible.

Our approach captures both the biophysical properties and the sequence context essential for accurate TF binding predictions. The surrogate model’s ability to generate features rapidly and at scale facilitates real-time, large-scale genomic analyses. These generated features are integrated into EPBDxDNABERT-2 (Kabir et al. 2024), a foundational genomic model that integrates DNA breathing, based on transformer architecture, to predict TF binding sites with enhanced accuracy. This integration not only improves prediction performance but also maintains the model’s interpretability, providing insights into how specific DNA sequences and their biophysical properties influence TF binding.

The implications of this work are significant for genomic research and biomedical applications. Rapid and accurate generation of DNA breathing features is essential for identifying new TFs, detecting regulatory mutations associated with diseases, and accelerating drug discovery efforts (Visscher et al. 2012). Our scalable, AI-driven solution opens new avenues for studying genetic variations, elucidating disease mechanisms, and developing targeted therapeutics.

In this paper, we detail the development and validation of our deep surrogate generative model for efficient DNA sequence feature generation. Our main contributions are summarized as below:

- We propose a novel conditional DDPM as a surrogate model to generate high-fidelity EPBD features conditioned on DNA sequences.
- We demonstrate that the surrogate model significantly reduces computational time from days to hours with minimal drop in performance.
- We integrate surrogate-generated features into the EPBDxDNABERT-2 model to enhance TF binding site predictions.
- We provide comprehensive evaluations showing that our approach performs comparably to traditional methods while offering substantial computational advantages.

## 2 Background and Related Work

Advancements in high-throughput sequencing technologies have expanded our ability to explore the human genome’s complexity (Shendure and Ji 2008). Modeling DNA dynamics, particularly DNA breathing, is crucial for elucidating how TFs access specific binding sites, impacting gene expression (Alexandrov et al. 2012; Nowak-Lovato et al. 2013b). Traditional methods like MCMC and Langevin dynamics simulate DNA breathing dynamics with high fidelity (Peyrard and Bishop 1989; Alexandrov et al. 2010), but are computationally intensive. Computational models have been developed to predict TF binding sites from DNA sequences. The early methods relied on position weight matrices (PWMs) (Stormo 2000), but lacked specificity. Deep learning models such as DeepBind (Alipanahi et al. 2015) and DeepSEA (Zhou and Troyanskaya 2015) use convolutional neural networks to learn sequence patterns associated with TF binding. Transformer-based architectures, such as DNABERT (Ji et al. 2021), model long-range dependencies in genomic sequences.

Recent advancements in generative modeling have significantly impacted genomic research, particularly in the synthesis and design of DNA sequences. Killoran et al. (Killoran et al. 2017) introduced deep generative models capable of creating synthetic DNA sequences, capturing essential structures for applications such as protein-binding microarrays. Building on this, Ho et al. (Ho, Jain, and Abbeel 2020) developed Denoising Diffusion Probabilistic Models (DDPMs), which have been applied to various domains, including genomics. Leveraging these models, DaSilva et al. (Senan et al. 2024) presented DNA-Diffusion, a conditional diffusion model designed to generate context-specific DNA regulatory sequences, demonstrating the potential of diffusion models in genomic sequence design. However, the existing works focused on sequence generation rather than generating biophysical features conditioned on sequences. Our work is distinct in utilizing a conditional DDPM to generate EPBD features, bridging the gap between sequence data and biophysical properties.

## 3 Methodology

Our surrogate model utilizes a conditional DDPM to generate EPBD features conditioned on DNA sequences. As illustrated in Figure 1, the model architecture consists of two primary components: a sequence embedder and a UNet-based denoising network. The overall pipeline comprises three main stages. i) DNA sequences and their corresponding EPBD features are obtained using pyDNA-EPBD simulations to provide the necessary sequence-feature pairs for training. ii) The surrogate model comprising the sequence embedder and the conditional DDPM, is trained to learn the mapping from DNA sequences to EPBD features. iii) The trained model is deployed to generate EPBD features for new DNA sequences. By inputting DNA sequences into the model, we efficiently obtain high-fidelity EPBD features without the computational overhead of traditional simulations.

**Figure 1:**
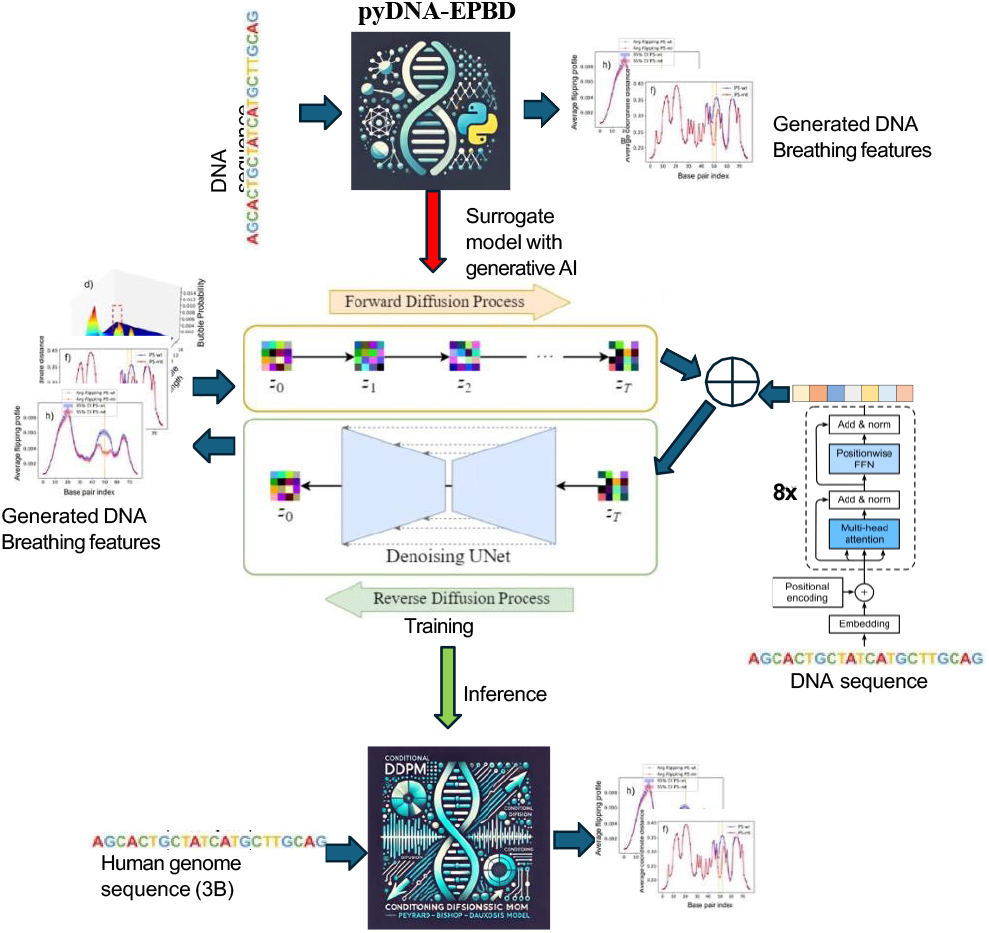
Overview of the surrogate model pipeline. (1) DNA sequences and corresponding EPBD features are obtained using pyDNA-EPBD simulations. (2) The surrogate model, consisting of a sequence embedder and a conditional DDPM, is trained on these pairs. (3) At inference, the model generates EPBD features conditioned on new DNA sequences.

### 3.1 Dataset

We constructed a dataset of DNA sequence and Extended Peyrard-Bishop-Dauxois (EPBD) feature pairs. The EPBD features were generated using biophysical simulations, specifically MCMC methods, capturing DNA breathing dynamics (Alexandrov et al. 2010). Due to computational intensity, we generated EPBD features for a subset of the genome, focusing on sequences of length *L* = 200 base pairs. The dataset is organized by partitioning the chromosomes into distinct subsets for training, validation, and testing to ensure robust model evaluation. Specifically, chromosomes 1 through 6, 10 through 22, designated as the training set, providing the primary data for model learning. Chromosome 7 serves as the validation set, enabling fine-tuning of hyperparameters and monitoring performance during training. Finally, chromosomes 8 and 9 are reserved exclusively as the test set, ensuring an unbiased evaluation of the model’s generalization capability on unseen data.

### 3.2 Sequence Embedder

The sequence embedder transforms DNA sequences into continuous embeddings that serve as conditioning information for the diffusion model. Given a DNA sequence **s** = *{s*_1_, *s*_2_, …, *s*_*L*_*}*, where each nucleotide *s*_*i*_ is represented by symbol from the set *{*A, C, G, T, N*}*, we map each nucleotide to a learnable embedding vector. Specifically, each nucleotide is first converted into a one-hot encoded vector and then projected into a continuous space using an embedding matrix 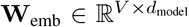, where *V* = 5 is the vocabulary size (including “N” for any nucleotide), and *d*_model_ is the embedding dimension:

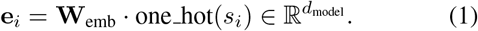

To incorporate positional information, we add sinusoidal positional encodings (Vaswani et al. 2017) to the embeddings:

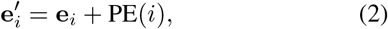

where PE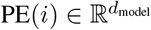 is defined as:

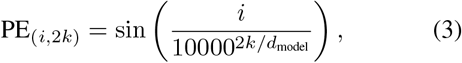

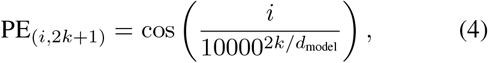

for *k* = 0, 1, …, *d*_model_*/*2 − 1.

The sequence of embeddings 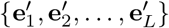 is then processed by a Transformer encoder (Vaswani et al. 2017), which consists of *N* layers of multi-head self-attention and position-wise feed-forward networks. The output of the Transformer encoder is a set of hidden states {**h**_1_, **h**_2_, …, **h**_*L*_}, where 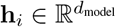.

To obtain a fixed-length context vector 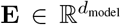 that summarizes the sequence information, we apply an attention pooling mechanism. We compute attention weights *α*_*i*_ for each position *i*:

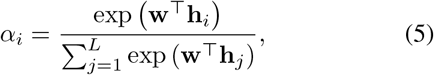

where 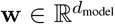 is a learnable parameter vector. The context vector **E** is then computed as a weighted sum of the hidden states:

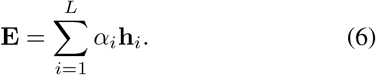

We set the embedding dimension to *d*_model_ = 256 and use *N* = 6 Transformer encoder layers. The multi-head self-attention mechanism employs 8 attention heads. These choices balance model capacity and computational efficiency.

### 3.3 Conditional Denoising Diffusion Probabilistic Model for EPBD feature generation

Our conditional DDPM models the conditional distribution *p*_*θ*_(**x**_0_ |**E**), where **x**_0_ is the EPBD feature corresponding to a DNA sequence, and **E** is the sequence embedding. The model learns to generate EPBD features by reversing a forward diffusion process, conditioned on **E**.

#### Forward Diffusion

In the forward diffusion process, starting from the original EPBD feature **x**_0_, Gaussian noise is progressively added over *T* timesteps to produce a sequence of noisy features {**x**_1_, **x**_2_, …, **x**_*T*_}. At each timestep *t*, the noisy EPBD feature **x**_*t*_ is obtained using the following equation:

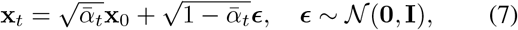

where 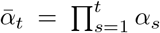, and 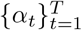 is a predefined noise schedule that determines the rate at which noise is added.

#### Reverse Diffusion

The reverse diffusion process aims to recover **x**_0_ from the noisy observation **x**_*T*_ by iteratively denoising through all timesteps in reverse order. At each timestep *t*, the model predicts the noise component ***ϵ***_*θ*_(**x**_*t*_, *t*, **E**) using the denoising network. The estimate of **x**_*t*−1_ is computed as:

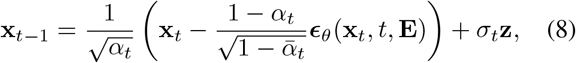

where *σ*_*t*_ is the standard deviation of the noise added at timestep *t*, and **z** ∼ 𝒩(**0, I**) if *t >* 1, or **z** = **0** if *t* = 1. The reverse process iteratively refines the estimate of the EPBD feature until it reaches **x**_0_.

#### UNet-based Denoising Network

The UNet-based denoising network operates on the EPBD feature space and models the reverse diffusion process. It is designed to predict the noise added to the EPBD features at each timestep, conditioned on the noisy input **x**_*t*_, the timestep *t*, and the sequence embedding **E**. The network architecture follows the UNet design (Ronneberger, Fischer, and Brox 2015), consisting of an encoder (downsampling path) and a decoder (upsampling path) with skip connections that concatenate feature maps from corresponding layers.

In our implementation, The UNet architecture consists of 4 downsampling and 4 upsampling blocks. Each block contains convolutional layers with kernel size 3, stride 2 for downsampling, and transposed convolutions for upsampling. We use ReLU activation functions and apply dropout with a rate of 0.1 to prevent overfitting.

#### Conditional Information Incorporation

Conditional information is integrated into the denoising network through adaptive normalization layers. Specifically, we use Adaptive Group Normalization (Dhariwal and Nichol 2021), where the scaling and bias parameters are modulated based on the sequence embedding **E** and the timestep embedding **t**_emb_. The timestep *t* is embedded using sinusoidal positional encodings:

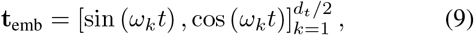

where 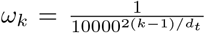, and *d*_*t*_ is the dimensionality of the timestep embedding. The combined conditioning vector is formed by integrating **E** and **t**_emb_, which is then used to modulate the normalization layers throughout the network.

#### Noise Schedule

We adopt a linear noise schedule where *α*_*t*_ decreases linearly from *α*_1_ = 1 − *β*_1_ to *α*_*T*_ = 1 − *β*_*T*_, with *β*_*t*_ linearly increasing from *β*_1_ = 1 × 10^−4^ to *β*_*T*_ = 0.02 over *T* = 1000 timesteps. This schedule balances the smoothness of noise addition and the stability of the reverse diffusion process.

#### Training Objective

The model is trained to minimize the expected mean squared error between the true noise ***ϵ*** added to **x**_0_ and the predicted noise ***ϵ***_*θ*_(**x**_*t*_, *t*, **E**):

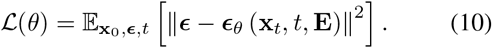

This objective function encourages the model to accurately predict the noise component at each timestep, which is critical for effective denoising during the reverse diffusion process.

### 3.4 Training Procedure

The training procedure involves optimizing the model parameters *θ* using stochastic gradient descent based on the training objective. For each iteration, a batch of DNA sequences and their corresponding EPBD features is sampled. The DNA sequences are processed through the sequence embedder to obtain the conditioning embeddings **E**. A random timestep *t* is sampled uniformly from {1, 2, …, *T*}, and Gaussian noise ***ϵ*** ∼ 𝒩(**0, I**) is generated.

The noisy EPBD feature **x**_*t*_ is computed using the forward diffusion equation. The denoising network then predicts the noise component ***ϵ***_*θ*_(**x**_*t*_, *t*, **E**). The loss is calculated as the mean squared error between the true noise ***ϵ*** and the predicted noise. The model parameters are updated using the AdamW optimizer (Loshchilov and Hutter 2019) with a learning rate of 8 *×* 10^−5^ and weight decay of 0.1. Gradient clipping with a maximum norm of 1.0 is applied to stabilize training. An Exponential Moving Average (EMA) of the model parameters is maintained with a decay rate of 0.995 to improve generalization.

We trained the surrogate model on a dataset comprising more than a million DNA sequence of length 200 bases and EPBD feature pairs. The sequences were randomly sampled from the human genome to capture a diverse range of genomic contexts. Training was conducted over 100 epochs with a batch size of 128. We employed a cosine annealing learning rate scheduler starting from 8 *×* 10^−5^ to enhance convergence.

### 3.5 Generation Process

After training, the model can generate EPBD features conditioned on new DNA sequences by performing the reverse diffusion process starting from pure noise. Given a DNA sequence **s**, we compute the sequence embedding **E** using the sequence embedder. We initialize **x**_*T*_ by sampling from a standard normal distribution **x**_*T*_∼𝒩 (**0, I**). Starting from *t* = *T*, we iteratively apply the denoising step:

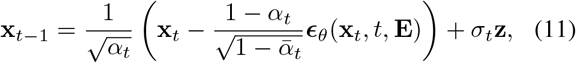

and sampling **z** ∼ 𝒩 (**0, I**) if *t >* 1. This process continues until *t* = 1, resulting in the generated EPBD feature **x**_0_ conditioned on the DNA sequence **s**.

### 3.6 Integration with EPBDxDNABERT-2

The surrogate-generated EPBD features are integrated into the EPBDxDNABERT-2 model (Kabir et al. 2024), which combines DNA sequence information with biophysical features for TF binding site prediction as shown in Figure 2. The EPBDxDNABERT-2 model consists of a DNABERT-2 encoder and a cross-attention mechanism to fuse the EPBD features with the sequence embeddings.

**Figure 2:**
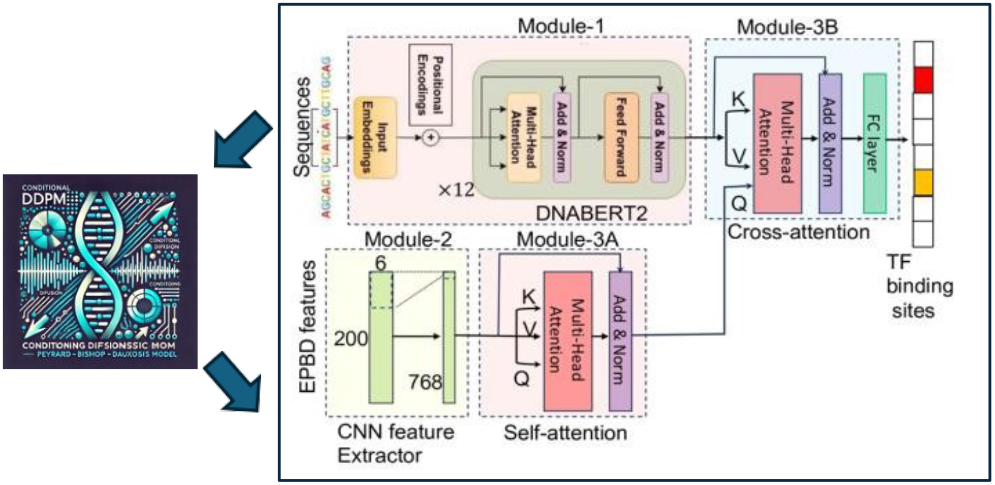
Demonstration of Transcription Factor Binding Prediction Using the Multimodal Foundational Model EPBDxDNABERT-2 with Features Generated by the Surrogate Model.

#### Evaluation

We evaluated the surrogate model’s performance and its impact on TF binding prediction using a heldout test set. The evaluation focused on two key aspects: the fidelity of the surrogate-generated EPBD features and the effectiveness of these features in downstream TF binding prediction tasks.

To assess the fidelity, we compared the surrogate-generated EPBD features with the ground truth features obtained from traditional biophysical simulations using the Mean Squared Error (MSE):

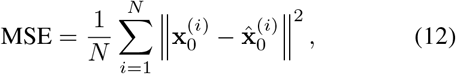

where *N* is the number of test samples, 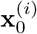 is the ground truth EPBD feature, and 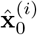 is the surrogate-generated feature.

For TF binding prediction, we integrated the surrogate-generated features into the EPBDxDNABERT-2 model and evaluated the model’s performance using the Area Under the Receiver Operating Characteristic Curve (AUROC) and the Area Under the Precision-Recall Curve (AUPR).

### 3.7 Implementation Details

Our implementation was developed using Python 3.11 and the PyTorch deep learning framework. PyTorch Lightning was employed to streamline the training process and manage multi-GPU support, which simplifies the codebase and allows for efficient experimentation. Training was conducted on high-performance computing clusters equipped with four NVIDIA GH200 GPUs per node, leveraging Distributed Data Parallel (DDP) training to maximize computational efficiency. All hyperparameters, such as learning rates, batch sizes, and network architectures, were selected based on hyperparamter tuning. We employed mixed-precision training to reduce memory consumption and accelerate computation without compromising numerical stability.

## 4 Results and Discussions

### 4.1 Surrogate Model Fidelity

To evaluate whether the diffusion process accurately learns the underlying distribution of features at each base position, we performed the Kolmogorov–Smirnov (KS) two-sample test on both the validation and test sets. Our hypothesis posits that the generated samples originate from the same distribution as the ground truth samples. We randomly sampled 1,000 examples from the validation and test sets and used our generative model to produce the corresponding EBPD features. For each base position, we computed the KS two-sample test, yielding test statistics and p-values for the 1,000 sampled examples at every base pair index. This analysis was repeated across 10 independent trials, and the mean p-values and test statistics were calculated for each feature at each base pair index.

Using a confidence level of 95%, we identified combinations of features and base pair indices where the average p-values were less than 0.05, indicating significant differences between the generated and ground truth distributions. These results were visualized in Figure 3, with subplots depicting the validation set (A) and test set (B). The figure reveals that, for most feature-index combinations, the p-values exceed 0.05, indicating that we cannot reject the null hypothesis in favor of the two-sided alternative. This suggests that the data generated by the diffusion process is statistically indistinguishable from the ground truth distribution for most features and base pair positions, demonstrating its potential to faithfully replicate the underlying patterns of the data. Regions highlighted in green represent combinations where the null hypothesis could not be rejected.

**Figure 3:**
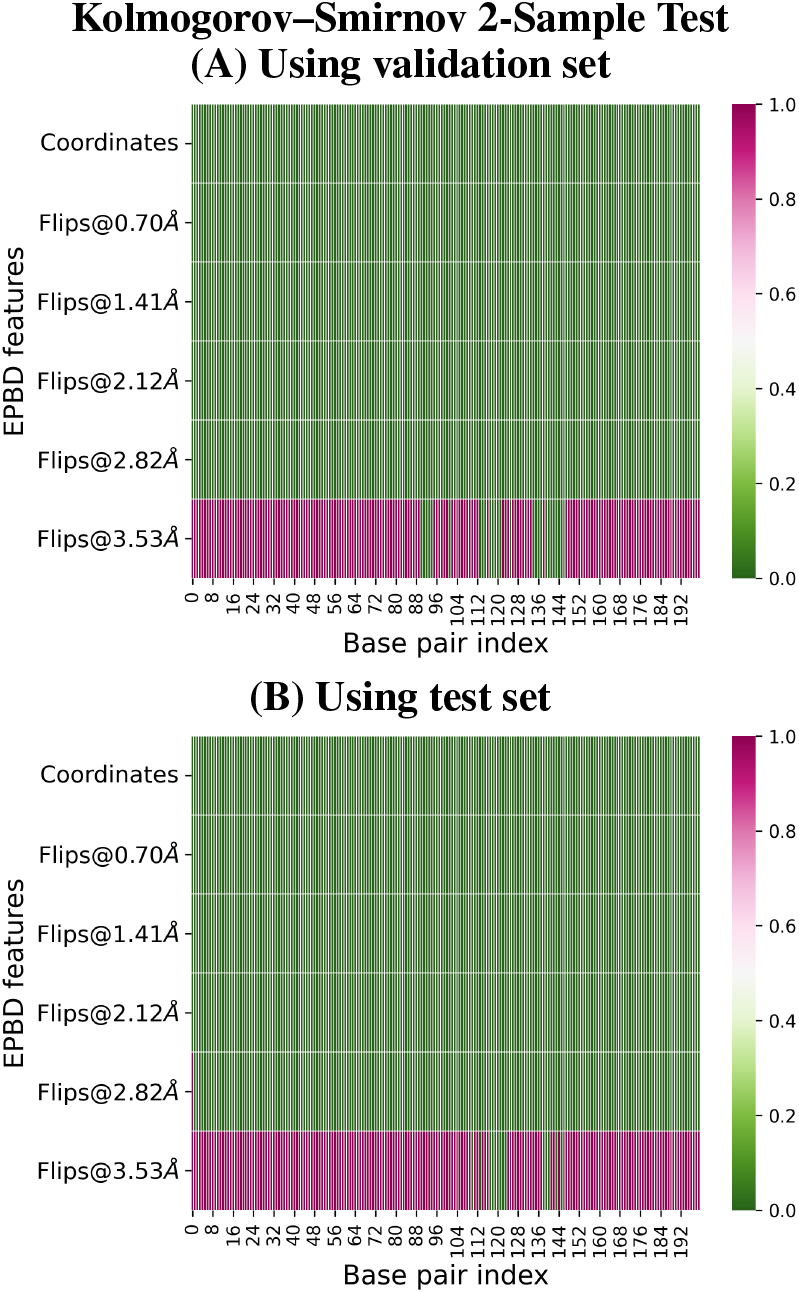
Visualization of the KS two-sample test results. Green regions indicate feature-base pair index combinations where the null hypothesis (that the surrogate-generated and ground truth samples come from the same distribution) cannot be rejected at the 95% confidence level. The results suggest that the surrogate model effectively learns the distribution of most features across base pair positions.

### 4.2 Feature Distribution Comparison

We evaluated the surrogate model’s performance by comparing its generated features with ground truth data for 10,000 sequences, achieving a low mean squared error (MSE) of 0.0025, which indicates high fidelity in replicating DNA dynamics. Furthermore, using thresholds ranging from 0.7071*Å* to 3.5355*Å*, we examined DNA breathing phenomena such as base flipping probabilities, a critical factor in processes like transcription factor binding and structural stability. The model’s ability to produce flipping profiles that closely align with the pyDNA-EPBD ground truth, particularly for wild-type and mutant promoter sequences of Adeno-Associated Virus (AAV) P5, as shown in Figure 4, demonstrates its ability to distinguish subtle structural differences critical for transcription regulation. The left and right panels provide a visual comparison between the generated features and the ground-truth features produced by EPBDSurr and pyDNA-EPBD, respectively. These panels illustrate the degree of similarity, highlighting how closely the generated features align with the expected ground-truth values.

**Figure 4:**
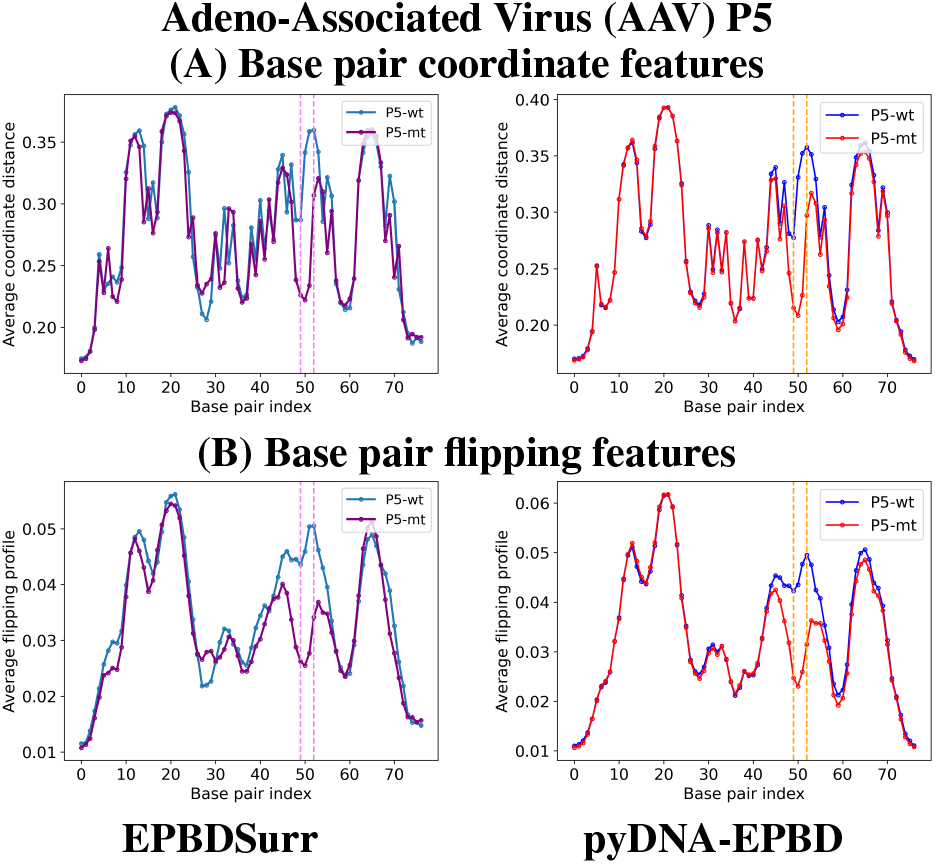
(A) Average coordinate distance and (B) flipping profile for AAV P5 wild- and mutant-promoter sequences at each base pair generated by EPBDSurr (left-panel) and pyDNA-EPBD (right-panel) model. The vertical blocks (purple (left-panel) and yellow (right-panel)) shows the mutation of a nucleotide bps substitutions from AG to TC at 50 and 51 positions (zero-indexed).

### 4.3 Transcription Factor Binding Prediction

We evaluated the impact of the surrogate-generated EPBD features on TF binding prediction using the EPBDxDNABERT-2 model. The performance was compared against baseline models and the model using ground truth EPBD features. The results are summarized in Table 1.

**Table 1:**
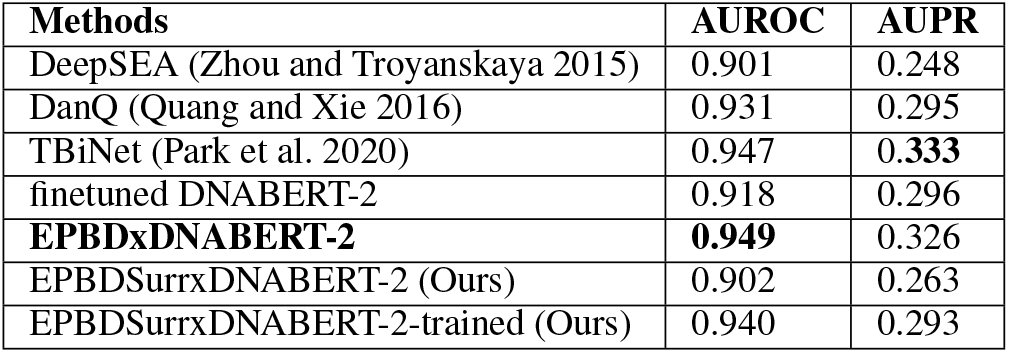
Overall performance comparison of our model (EPBDSurrxDNABERT2) with baseline works in light of AUROC and AUPR metrics.

| Methods | AUROC | AUPR |
| --- | --- | --- |
| DeepSEA (Zhou and Troyanskaya 2015) | 0.901 | 0.248 |
| DanQ (Quang and Xie 2016) | 0.931 | 0.295 |
| TBiNet (Park et al. 2020) | 0.947 | **0.333** |
| finetuned DNABERT-2 | 0.918 | 0.296 |
| **EPBDxDNABERT-2** | **0.949** | 0.326 |
| EPBDSurrxDNABERT-2 (Ours) | 0.902 | 0.263 |
| EPBDSurrxDNABERT-2-trained (Ours) | 0.940 | 0.293 |

The model using surrogate-generated features without fine-tuning (EPBDSurrxDNABERT-2) achieved an AUROC of 0.902 and an AUPR of 0.263, demonstrating that even without retraining, the surrogate features provide valuable information for TF binding prediction. After fine-tuning the EPBDxDNABERT-2 model with surrogate-generated features (EPBDSurrxDNABERT-2-Trained), the AUROC improved to 0.940 and the AUPR to 0.293, closely approaching the performance of the model using ground truth EPBD features. These results highlight the surrogate model’s effectiveness in capturing essential biophysical characteristics necessary for accurate TF binding site prediction.

## 5 Runtime and Speedup Analysis for pyDNA-EPBD and EPBDSurr

The runtime comparison between pyDNA-EPBD and EPBDSurr for CPU/GPU clusters for different batch sizes is presented in Table 2 and Figure 5. This study was conducted in a cluster where each node has 2 AMD EPYC 7713 Processors and 4 NVIDIA Ampere A100 GPUs. The AMD EPYC 7713 CPUs have 64 cores peaking at 3.67 GHz and 256 GB RAM. For a batch size of 256, the pyDNA-EPBD executed in CPU cluster requires 52.4 minutes, whereas the EPBDSurr in GPU cluster completes the same workload in just 0.3058 minutes, yielding a speedup of approximately 172x. As the batch size increases, EPBDSurr runtimes scale more efficiently compared to the linear increase observed in pyDNA-EPBD runtimes. At the largest batch size of 8192, the pyDNA-EPBD requires 1,728 minutes, whereas the EPBDSurr cluster completes the workload in 8.3 minutes, resulting in a speedup of 208.43x. These results demonstrate the significant computational advantage of EPBDSurr in leveraging GPUs for handling large-scale data processing tasks.

**Table 2:** Runtime in minutes and speedup analysis for EPBDSurr over pyDNA-EPBD across different batch sizes.

| Batch Size | pyDNA-EPBD | EPBDSurr | Speedup |
| --- | --- | --- | --- |
| 256 | 52.4 | 0.305 | 171.80 |
| 512 | 108 | 0.59 | 183.05 |
| 1,024 | 216 | 1.1 | 196.36 |
| 2,048 | 432 | 2.1 | 205.71 |
| 4,096 | 864 | 4.2 | 205.71 |
| 8,192 | 1,728 | 8.3 | 208.43 |

**Figure 5:**
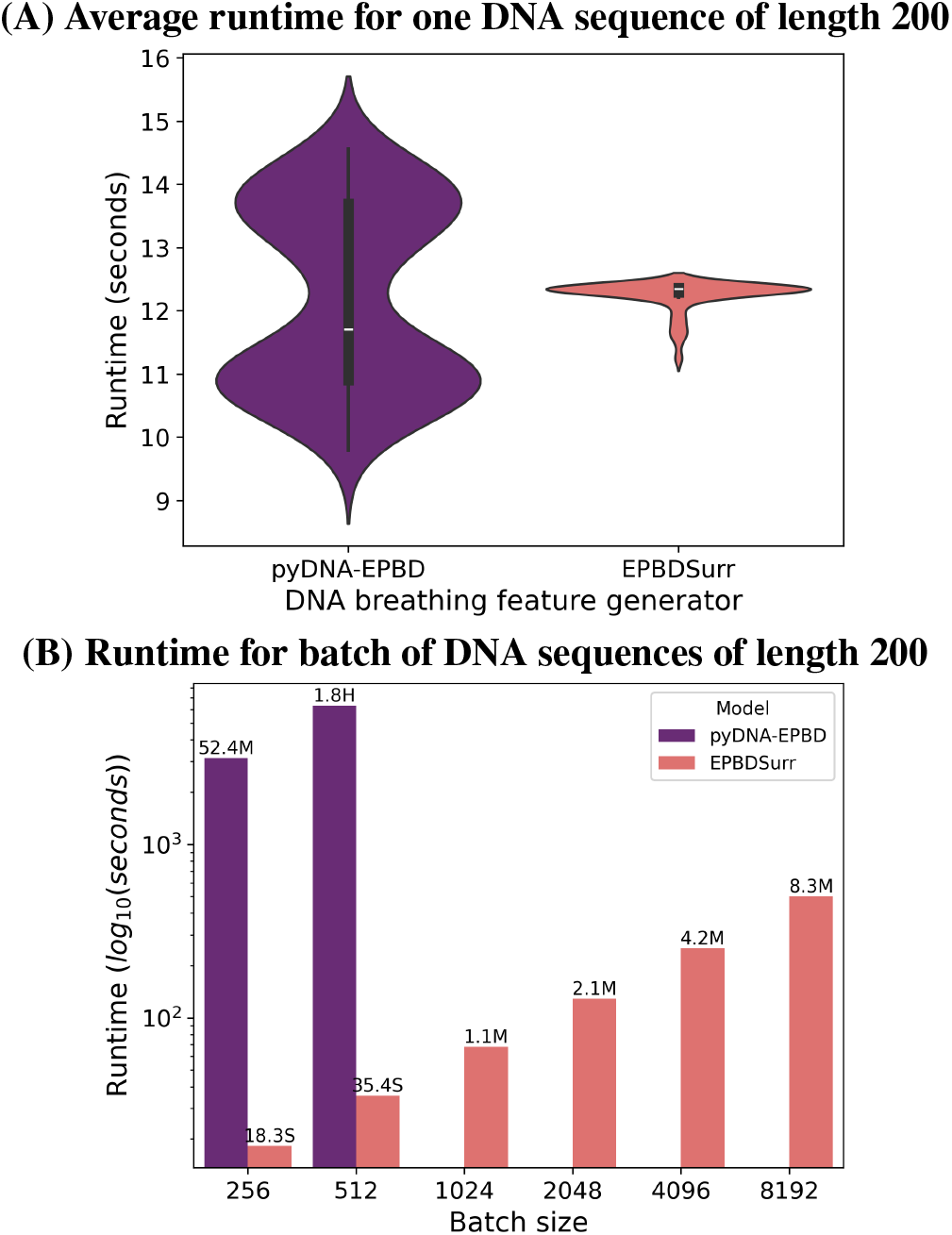
(A) Average runtime for generating DNA breathing features for one DNA sequence of length 200 nucleotide base pairs. (B) Runtime for generating DNA breathing features for batch of sequences.

The runtime comparison between pyDNA-EPBD and EPBDSurr for large genomic workloads further underscores the utility of GPUs for scaling data-intensive pipelines. For ChIP-seq data comprising 3 million sequences, the pyDNA-EPBD running on a CPU cluster with 100 nodes, each processing a batch of 8,192 sequences in 1,728 minutes, takes approximately 115.2 hours (4.8 days). In contrast, EPBDSurr leveraging a GPU cluster with 100 nodes, each equipped with 4 A100 GPUs processing sequences in batches of 8,192, completes the same workload in just 8.3 minutes. Similarly, for the human genome, which comprises 3 billion sequences, the pyDNA-EPBD cluster requires an impractical 105,486 hours (approximately 4,395 days), whereas the EPBDSurr completes the task in a mere 126.88 hours (approx 5.29 days) on an A100 GPU cluster, or approximately 63.44 hours (approx 2.64 days) on an H100 GPU cluster with 100 nodes. This immense speedup—approx 833x for ChIP-seq data and 832x to 1,663x for the human genome—highlights the scalability and efficiency of EPBDSurr over pyDNA-EPBD.

## Supporting information

Overview of framework

## 6 Limitations

While our approach significantly reduces computational overhead, potential approximation errors introduced by the surrogate model may affect downstream analyses. Despite the generally promising results, we observed a notable exception: the diffusion process consistently failed to accurately generate flipping features associated with a 3.53 Å distance. This limitation is particularly striking, as flipping features are critical indicators of specific molecular interactions and structural conformations. The corresponding feature values at the 3.53 Å distance may have inherently low signal strength, making them difficult for the generative model to capture. These features might have a smaller magnitude relative to other features, leading the model to prioritize features with more pronounced signals during the learning process. This also highlight potential limitations in the model’s ability to capture complex or sequence-specific DNA behaviors, such as non-canonical structures or localized melting. These mismatches could impact predictions of biologically significant events, such as transcription efficiency or protein-binding site accessibility. While the surrogate model is computationally efficient and largely accurate, further refinement—such as expanding training data diversity and incorporating environmental factors—would enhance its generalizability and reliability, ensuring robust performance across diverse genomic contexts.

## 7 Conclusion

We have presented a deep surrogate generative model leveraging a conditional DDPM to efficiently generate high-fidelity DNA breathing features conditioned on DNA sequences. By integrating these features into the EPBDxDNABERT-2 model, we achieved comparative performance in TF-DNA binding site prediction accuracy while dramatically reducing computational overhead from months to hours. This advancement enables large-scale genomic analyses previously hindered by computational constraints, with significant implications for understanding gene expression regulation, disease mechanisms, and the development of precision medicine approaches.

## 8 Acknowledgements

This research project was supported by the LDRD 20230287ER (to M.B.), National Institute of Health RO1HL128831 (to A.U.) and National Institute of Health 5R01MH116281-03 (to B.A.). A.K. and A.S. were supported in part by the National Science Foundation Grant No. 2310113. This research used resources provided by the Los Alamos National Laboratory Institutional Computing Program, which is supported by the U.S. Department of Energy National Nuclear Security Administration under Contract No. 89233218CNA000001.

## Notes

### Competing Interest Statement

The authors have declared no competing interest.

