## Supplementary figures and images for "Scalable DNA Feature Generation and Transcription Factor Binding Prediction via Deep Surrogate Models"

### Overview of framework

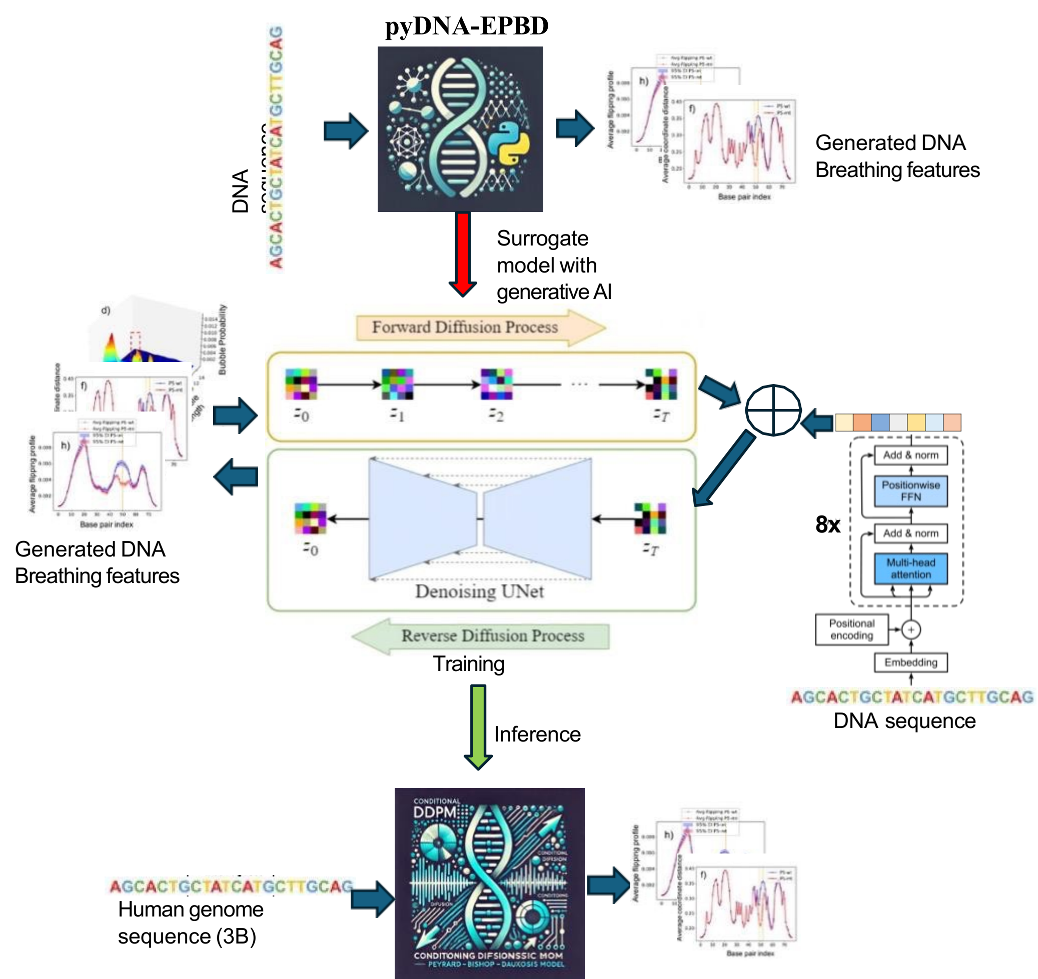
